# Infrared Nanospectroscopy Reveals the Molecular Interaction Fingerprint of an Aggregation Inhibitor with Single Aβ42 Oligomers

**DOI:** 10.1101/2020.06.24.168997

**Authors:** Francesco Simone Ruggeri, Johnny Habchi, Sean Chia, Michele Vendruscolo, Tuomas P. J. Knowles

## Abstract

Very significant efforts have been devoted in the last twenty years to developing compounds that can interfere with the aggregation pathways of proteins related to misfolding disorders, including Alzheimer’s and Parkinson’s diseases. However, no disease-modifying drug has become available for clinical use to date for these conditions. One of the main reasons for this failure is the incomplete knowledge of the molecular mechanisms underlying the process by which small molecules interact with protein aggregates and interfere with their aggregation pathways. Here, we leverage the single molecule level morphological and chemical sensitivity of infrared nanospectroscopy to provide the first direct measurement of the interaction between single Aβ42 oligomeric and fibrillar species and an aggregation inhibitor, bexarotene, originally an anticancer drug capable recently shown to be able to inhibit Aβ42 aggregation in animal models of Alzheimer’s disease. Our results demonstrate that the carbonyl group of this compound interacts with Aβ42 aggregates through a single hydrogen bond. These results establish infrared nanospectroscopy as powerful tool in structure-based drug discovery for protein misfolding diseases.

---

Alzheimer’s disease (AD) is characterised by memory loss and cognitive impairment. AD is the primary cause of dementia, which affects currently over 50 million people worldwide, a number expected to exceed 150 million by 2050.^1-3^ The self-assembly of the 42-residue isoform of the amyloid-β (Aβ) peptide into intractable aggregates is considered to be at the core of the molecular pathways leading to AD.^3-5^ It is therefore important to understand at the molecular level the aggregation process of Aβ42 in order to develop effective therapeutic strategies that aim at inhibiting its self-assembly.

Great efforts have been devoted in the last twenty years to understand the molecular basis of this devastating disorder and to develop small molecules that could interfere with the aggregation pathway of Aβ42.^6-12^ Indeed, disease-modifying small molecules represent c.a. one third of the registered trials, in which anti-Aβ therapies dominate.^13^ However, despite these efforts, no compound has entered the clinical use to date.^13,14^

These repeated failures are in part related to an incomplete understanding of the molecular mechanisms underlying the process by which small molecules interact with protein aggregates and how they interfere with the pathways of aggregation. It is increasingly recognised that inhibiting Aβ aggregation *per se* could have unexpected consequences on the toxicity, as it could not only decrease it, but also leave it unaffected, or even increase it.^15^ This complexity is due to the non-linear nature of the aggregation network, in which neurotoxic small oligomeric intermediates^16-20^, can be formed in a variety of ways, some of which highly sensitive to small perturbation. Therefore, promising effective therapeutic strategies must be aimed at targeting precise species in a controlled intervention during the aggregation process.

We have recently described a drug discovery strategy based on the accurate characterisation of the effects of small molecules on the kinetics of Aβ aggregation.^21^ This strategy exploits the power of chemical kinetics,^22,23^ in which the effect of small molecules on the rates of specific microscopic steps in Aβ42 aggregation can be determined quantitatively. Using this strategy, we have identified an FDA-approved anti-cancer drug, bexarotene, which was found to delay substantially the aggregation of Aβ42. Bexarotene delays the aggregation of Aβ42 in a concentration-dependent manner by inhibiting primary and secondary pathways.^21^ Although a kinetic understanding of the molecular mechanism by which small molecules affect Aβ aggregation has been achieved, the chemical interactions and the structural information on Aβ species in the free and bound forms to the small molecule remain to be unravelled. This limitation is mainly due to the lack of experimental tools that allow accurate direct measurements of the chemical and structural properties of individual species in a heterogeneous population.

A breakthrough in the analysis of the chemical properties of heterogeneous protein aggregation at the nanoscale has been achieved with the development and application of atomic force microscopy in combination with infrared nanospectroscopy (AFM-IR, **Fig. 1c**).^24-34^ AFM-IR is becoming widely applicable in biology since is capable of acquiring simultaneously morphological, nanomechanical and nanoscale-resolved chemical IR maps and absorption spectra from protein aggregates, liquid-liquid phase separated condensates, chromosomes, and single cells.^29,31,34-36^ As fundamental achievement, we have recently demonstrated that AFM-IR enables to acquire infrared absorption spectra for secondary structure determination from single protein molecules.^37^

**Figure 1.**
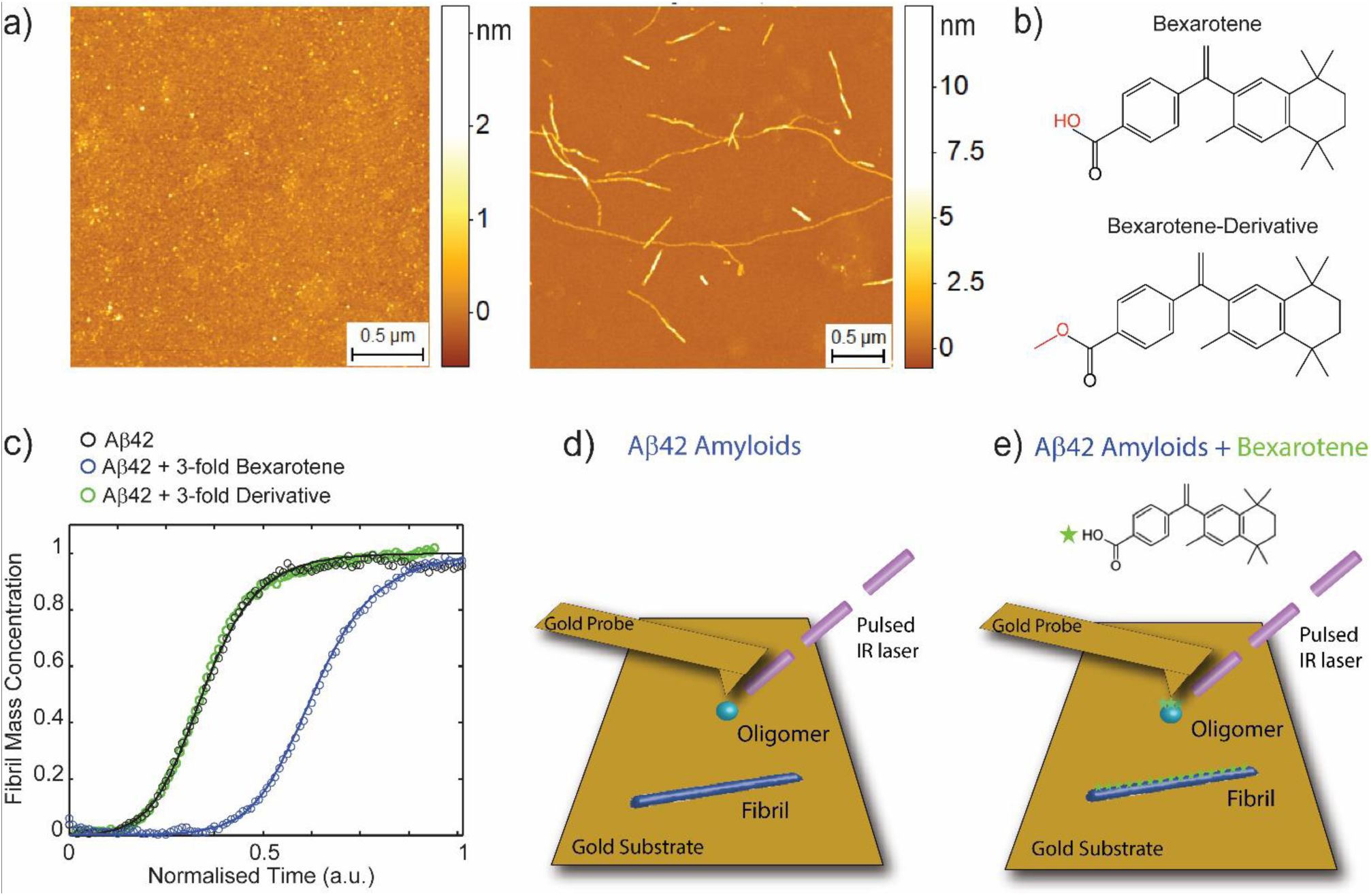
Effect of bexarotene on the aggregation kinetics of Aβ42. (a) Representative high resolution AFM morphology maps of Aβ42 aggregates before incubation and at the end of the aggregation reaction. (b) Structures of bexarotene and its derivative, where the hydrogen bond donor is replaced with a methyl group (highlighted in red). (c) Kinetics profiles of the aggregation reaction of 2 μM Aβ42 in the absence or in the presence of a 3:1 concentration ratio of bexarotene and its derivative. The solid lines show the best fit of the experimental data (open circles) when primary and secondary pathways are both inhibited by bexarotene. (c-d) The AFM-IR experimental approach to study the interaction of Aβ42 fibrils with bexarotene at the single aggregate scale, to reach high sensitivity the nanogap between a gold coated probe and a gold substrate is used.

In the present work, we exploit the powerful combination of single molecule imaging and vibrational spectroscopy offered by AFM-IR to characterise at the single aggregate scale the morphological, chemical and structural properties of the aggregate species of Aβ42 in the absence and the presence of bexarotene. We thus provide a direct measurement at the single oligomeric scale of the interactions between a small molecule and Aβ42 oligomers and fibrils. Our results constitute a key finding to investigate the transient and heterogeneous conformational changes occurring during protein aggregation, as well as the capability to study the interactions between proteins and therapeutic compounds, with the aim of developing therapeutics against neurodegenerative disorders.

## RESULTS

### Bexarotene Inhibits Aβ42 Aggregation

Before investigating the chemical interactions between bexarotene and Aβ42 aggregates at the nanoscale, we carried out a characterisation of the aggregation process of Aβ42 (**Fig. 1**). As shown by phase controlled high-resolution AFM imaging, before incubation at 0 h, Aβ42 is largely in a monomeric state and there are no fibrillar aggregates visible (**Fig. 1a**). After 4 h incubation, as previously reported, we observe an abundance of fibrillar aggregates with typical cross-sectional diameters between 2-6 nm (**Fig. S1**). We further performed a kinetic analysis of the aggregation of Aβ42 by a highly reproducible ThT-based protocol.^38^ In particular, we investigated the aggregation reaction in the absence and the presence of 3-fold excess of bexarotene and one of its derivatives where the carboxylic acid group was substituted by an ester functional group, leading to the absence of an acidic ionisable group (**Fig. 1c** and **S2-3**). The aggregation reaction displays the three phases of a typical nucleation-polymerisation reaction: lag, growth, and plateau. Indeed, typical amyloid formation displays sigmoidal growth kinetics, where a transition zone, namely the growth phase, is preceded and followed by flat regions, usually referred to as the lag phase and the plateau, respectively. Bexarotene delayed the aggregation of Aβ42 significantly, by inhibiting both primary and surface-catalysed secondary nucleation as previously demonstrated.^21^ On the other hand, its derivative did not affect the aggregation reaction even at high concentrations. These results show that bexarotene can slow down the kinetics of aggregation reaction through interaction with Aβ42 aggregates.

### Infrared Nanospectroscopy of Single Aβ42 Oligomers and Fibrils

We first applied conventional bulk infrared (IR) spectroscopy to investigate the chemical binding between Aβ42 fibrils formed at the end of the aggregation reaction (plateau phase) and bexarotene (**Fig. S4**). However, with bulk infrared spectroscopy we were not able to observe any difference between the fibrils formed in presence and absence of bexarotene, and thus we could not study in this way their chemical binding. This limitation is largely related to the nature of this bulk technique, which is not able to discriminate between the bound small molecules and the ones diffused in solution.

To overcome these limitations, we next exploited the nanoscale chemical resolution power of AFM-IR to study the chemical and structural properties of Aβ42 aggregates at the single molecule scale (**Fig. 1d-e**) in absence and presence of bexarotene. Consequently, ThT-free aliquots of Aβ42 solutions were removed at different time points from the aggregation reactions for single molecule measurements. In particular, we focused on the growth phase (2 h for Aβ42 alone and 4.5 h for Aβ42+bexarotene) and on the plateau phase of aggregation after 15 h incubation. We deposited the aliquots of Aβ42 aggregates on atomically flat (0.4 nm RMS) template gold surfaces. As expected, in the growth phase we observed the co-existence of oligomeric and fibrillar aggregates, while in the plateau phase we observed only fibrillar aggregates (**Fig. 2a-b** and **Fig. S5**), similarly as observed for samples deposited on conventional mica substrates (**Fig. 1**).

**Figure 2.**
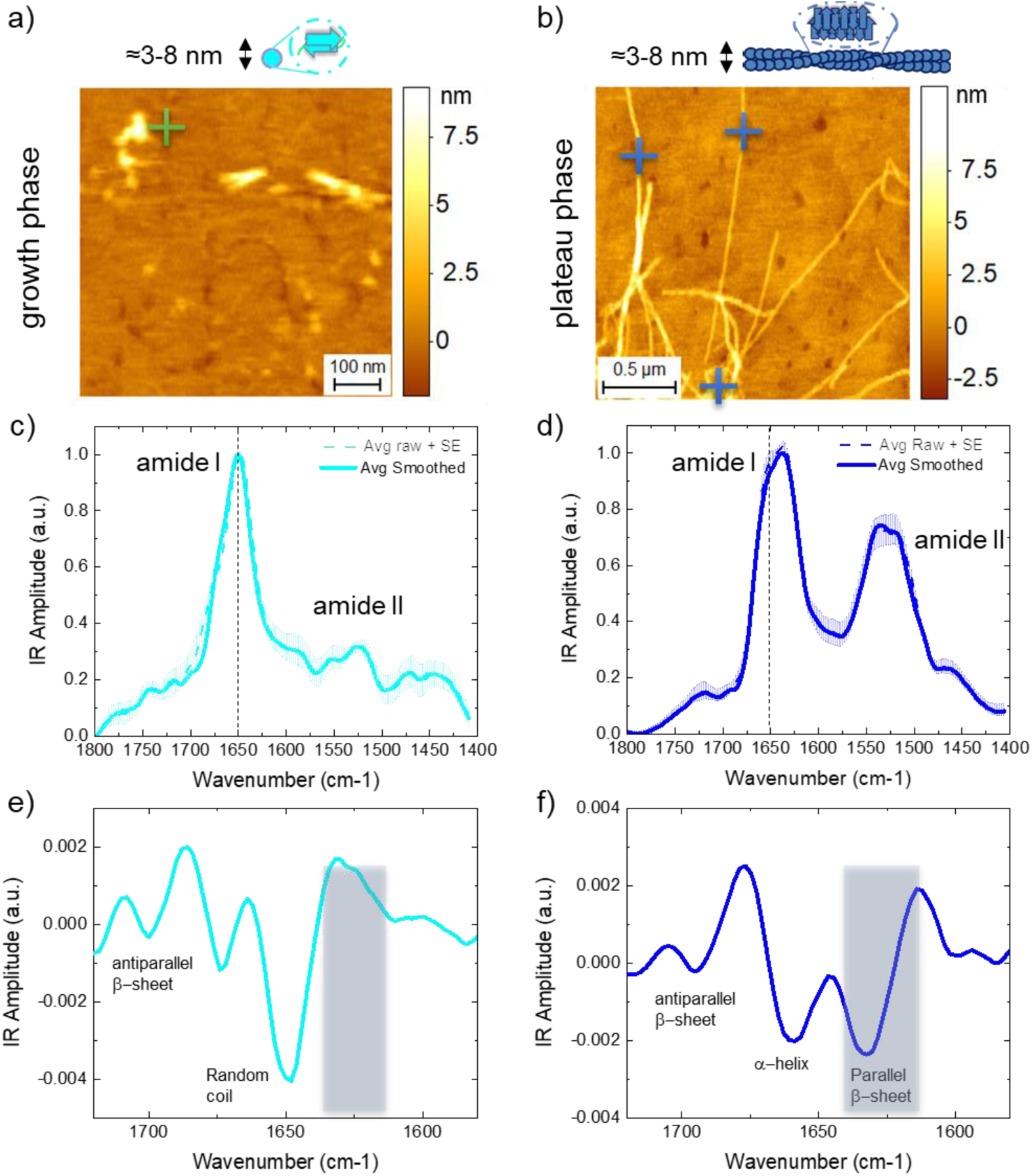
Single aggregate measurements of secondary and quaternary structure of Aβ42 aggregates. **(a**,**b)** AFM-IR morphology maps of oligomeric (a) and fibrillar species (b) of Aβ42. **(c**,**d)** Nanoscale localized IR absorption spectra of oligomeric (c) and fibrillar species (d) of Aβ42. We represent two spectra: i) the average + SE of the raw spectra, and ii) the average + SE of the preprocessed spectra by applying an adjacent averaging filter (5 pts) and a Savitzky-Golay filter (2^nd^ order, 15 pts). At least 5 different raw spectra at each positon (colored crosses) were acquired. **(e-f)** The second derivatives of the spectra are calculated with a Savitzky-Golay filter (2^nd^ order, 15 pts) to deconvolve the major secondary structural contributions of the oligomeric and fibrillar species.

Then, we focused on the measurement of the single aggregate chemical and structural properties of Aβ42 aggregates alone. The sensitivity of AFM-IR has been recently demonstrated to be capable to reach the detection of single protein molecules of apoferritin with a molecular weight of ∼400 kDa and hydrodynamic radius of ∼12 nm, with high signal-to-noise ratio in order to determine their secondary structure.^37^ This high sensitivity was reached by measuring at a short laser pulse, off resonance (ORS-nanoIR)^37^ and exploiting the field enhancement at the nanogap with the metallic probe and substrate and reach single aggregate sensitivity. We applied similar principles in the present study to reach the detection of the chemical signature of individual oligomeric (**Fig. 2c**) and fibrillar species of Aβ42 (**Fig. 2d**) with a typical cross-sectional diameter of both oligomeric and fibrillar ranging between ∼3-8 nm (**Fig. S6**). To exclude from our chemical and structural analysis the residual absorption of contaminants on the tip and on the substrate, we subtracted each spectrum acquired on one aggregate from the residual absorption on the substrate (**Fig. S7**).

We then applied second derivative analysis to deconvolve the amide band I of the aggregates and evaluate the structural contributions to their secondary and quaternary structure (**Fig. 2e-f**). The oligomeric aggregates presented their major structural contribution at 1648 cm^-1^, indicating the large presence of random coil conformation, but also a significant peak at 1695 cm^-1^ indicating the presence of intermolecular antiparallel β-sheets (**Fig. 2e**). In the case of the fibrillar aggregates, we could remarkably observe the presence of both the bands associated to a significant content of intermolecular parallel (1630 cm^-1^) and antiparallel β-sheet (1695 cm^-1^) (**Fig. 2f**). These results are in excellent agreement and demonstrate at the single aggregate scale the conversion step from an antiparallel β-sheet conformation in oligomeric species to a parallel β-sheet conformation in the fibrillar products of the aggregation kinetics.^39^ In the case of the oligomers, their cross-sectional diameter was below the size of a single protein molecule of apoferritin, which was only very recently achieved. Although we could reach this high sensitivity, the measurement of the spectra of the oligomers was at the limit of the experimental sensitivity leading to a higher experimental noise and a partial suppression of the amide band II.

### Interaction of Single Oligomeric and Fibrillar Species with Bexarotene

We next applied AFM-IR to unravel the chemical interaction and effect of bexarotene on the aggregated species of Aβ (**Fig. 3**). We acquired first AFM morphology maps of single oligomeric (**Fig. 3a**) and fibrillar (**Fig. 3b**) species formed during the kinetics of aggregation in the presence of bexarotene. The aggregates showed similar morphology and cross-sectional dimensions as the ones in absence of the small molecule (**Fig. S5**). Then, we acquired nanoscale resolved spectra from both oligomeric and fibrillar species incubated in presence of bexarotene (**Fig. 3c-d**). The spectra of the aggregates in presence of bexarotene showed marked differences when compared to the spectra of the aggregates incubated in absence of the small molecule. In particular, we observed a significantly increased absorption between 1750-1700 cm^-1^ and 1425-1475 cm^-1^ (green boxes in **Fig. 3c**,**d**). In order to prove that the spectral differences measured between aggregated species in presence and absence of bexarotene were statistically significant, we performed a principal component analysis (PCA, **Fig. 3e-g**).

**Figure 3.**
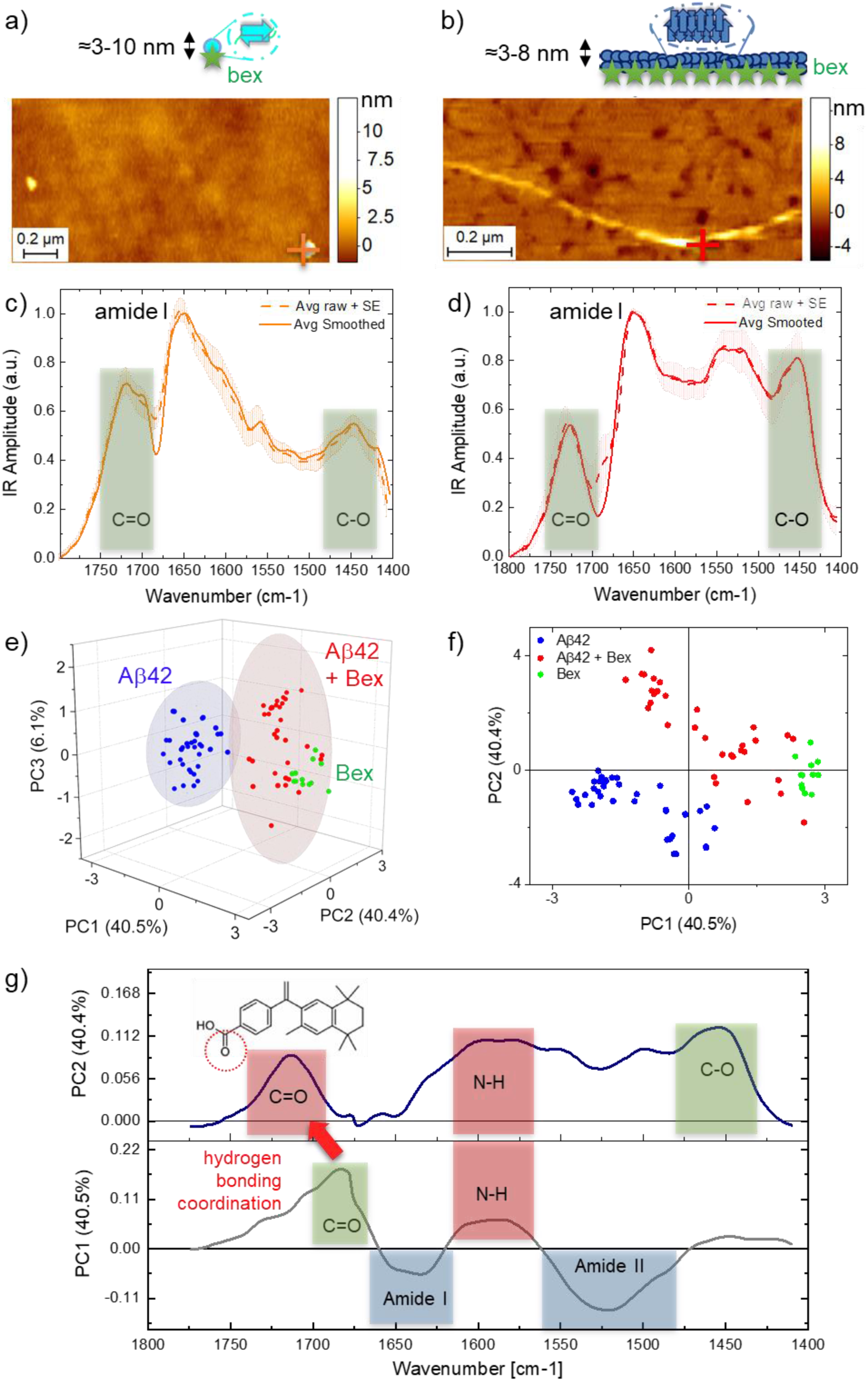
Structural characterisation of the interaction of bexarotene with single Aβ42 oligomers and fibrils. **(a-d)** AFM-IR morphology map (a,b) and nanoscale localized IR absorption spectra (c,d) of oligomeric (a,c) and fibrillar species (b,d) of Aβ42 incubated with bexarotene. We represent two spectra: i) the average +SE of the raw spectra, and ii) the average + SE of the preprocessed spectra by applying an adjacent averaging filter (5 pts) and a Savitzky-Golay filter (2^nd^ order, 15 pts). At least 5 different raw spectra at each positon (colored crosses) were acquired. **(e-f)** 3-D and 2-D score plots of PCA analysis applied to Aβ42 aggregates (oligomers and fibrils, n = 40) in absence of bexarotene (blue), Aβ42 aggregates in presence of bexarotene (oligomers and fibrils, n = 31) and bexarotene alone (n=9). The colored ellipsoids represent the 90% confidence of the variance of each groups, thus demonstrating the separation of the spectroscopic signature of the aggregates in presence and absence of the small molecule. **(g)** Loading plots of the PC1 and PC2.

PCA allowed noise reduction and the detection of subgroupings within the spectra of the aggregates in presence and absence of bexarotene. The score plots of the analysis (**Fig. 3e, f**) represent each single nanoscale resolved IR spectrum as a point in the multidimensional space of principal components (PCs). The first three PCs were sufficient to represent more than 85% of the spectra variability in the ensemble and demonstrate the high statistical significance of the sub-grouping (**Fig. 3e**) of the spectroscopic signature of the aggregates formed in absence (blue) and presence (red) of bexarotene. The spectral differences represented by each principal component are shown in the loading plots (**Fig. 4g and Fig. S8**), which show the spectroscopic region responsible for the greatest degree of separation inside between the spectra sub-groups. The first two PCs were sufficient to generalize the spectral differences (>80% spectral difference). A negative PC1 and PC2 were correlated with the chemical signature of the characteristic amide bands I (1700-1600 cm^-1^) and II (1580-1500 cm^-1^) of protein, dominating the spectra in absence of bexarotene. While positive PC1 and PC2 were correlated with the chemical signature of bexarotene at 1700-1750 cm^-1^ (C=O), 1580 cm^-1^ and 1450 cm^-1^ (C-O) (**Fig. 4a**). As expected, the spectra of the aggregates in presence of bexarotene (red) had mixed grouping properties when compared with the IR absorption signature of the small molecule (green) and the aggregates alone (blue). Thus, the analysis demonstrates with high statistical significance the presence of the chemical signature of bexarotene within the Aβ42 aggregates incubated with the molecule.

**Figure 4.**
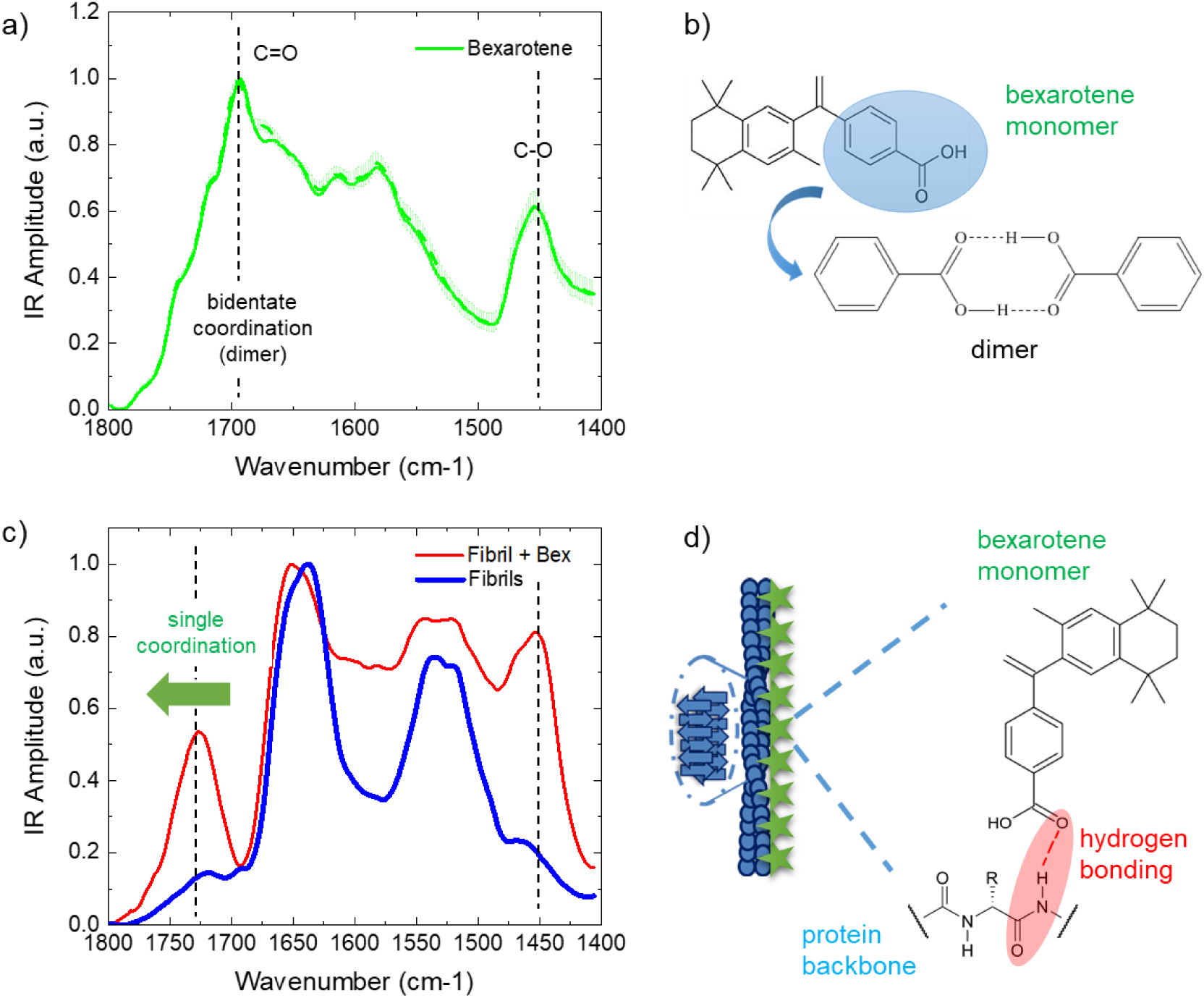
Bexarotene interacts with Aβ42 aggregates through hydrogen bonding of its carbonyl group. **(a)** AFM-IR spectrum of bexarotene molecule film deposited on gold substrate. **(b)** Bexarotene alone has strong tendency to form dimers with C=O IR absorption at 1690 cm^-1^, while in its monomeric form the drug has C=O IR absorption at 1725 cm^-1^. **(c)** Aβ42 fibrils show the typical spectrum of proteins (blue), while fibrils incubated with the drug show new chemical signature related to the drug in the monomeric for and interacting with the fibrils through hydrogen bonding (red). **(d)** Schematics of fibril-small molecule interaction. When bonded with the protein, bexarotene is likely to assume a conformation such that the bulky part of the molecule is projected away from the nearest amino acid side chains.

### Bexarotene Interacts with Aβ42 Aggregates through Hydrogen Bonding of its Carbonyl Group

The increased IR absorption in the spectra of the aggregates incubated with bexarotene showed two main peaks of absorption at 1750-1700 cm^-1^ and 1450 cm^-1^, related to the vibration of C=O and C-O bonds of the small molecule alone (**Fig. 4a**). Further fundamental differences between the spectrum of bexarotene and the aggregates in presence of the molecule were found. In particular, the loading plots showed a significant shift of the frequency of absorption of the carbonyl band of bexarotene when in presence of the amyloid aggregates (**Fig. 4g**). Bexarotene alone in solution has strong tendency to be in a dimeric form through a bidentate coordination (**Fig. 4b**), thus showing a typical frequency of the C=O bond at approximately 1690 cm^-1^ as previously shown in literature in the similar case of benzoic acid and its derivatives.^40^ Importantly, as also shown in the average spectra in **Fig. 4c** and in the loading plots of PCA (**Fig. 4g** and **S8**), the C=O peak of bexarotene in the presence of the aggregates is significantly shifted at higher wavenumbers (1725 cm^-1^). This chemical shift demonstrates that bexarotene is transitioning from a double coordination (dimeric form) to a monomeric form coordinating with the amyloid aggregates through only one hydrogen bonding, thus causing the shift of the carbonyl peak at higher energy (**Fig. S9**).^41^ This difference is also demonstrated by comparing the experimental spectrum of the fibrillar aggregates incubated with bexarotene and the calculated spectrum obtained summing the spectroscopic response of the fibrils and the drug alone (**Fig. S9**). The two spectra differ significantly in the region of the carbonyl group and of the N-H bond, which is involved in hydrogen bonding and thus demonstrating that this chemical bond is indeed involved in the protein-small molecule interaction. Other possible interactions between the carboxylic group and the protein are disfavoured due to molecular bulkiness, steric hindrance, and the relatively long distance between two consecutive parallel amide hydrogens in the polypeptide chain.

In summary, the analysis of the spectra demonstrates that hydrophilic moiety of bexarotene binds to Aβ42 mainly through a hydrogen bond to the proteins aggregates (**Fig. 4d**), in excellent agreement with the finding that the bexarotene derivative that does not contain the acidic ionisable group (hydrogen bond donor) is not capable of modifying the aggregation kinetics (**Fig. 1c**).

## DISCUSSION

In this study, we applied infrared nanospectroscopy in combination with multivariate data analysis to first study the secondary structure of Aβ42 oligomeric and fibrillar amyloid species at the single aggregate scale, and then prove their chemical interactions with a therapeutic compound with single bond resolution. Initially, we characterised the aggregates of Aβ42 proving a transition from antiparallel to parallel intermolecular β-sheet content when going from the oligomeric to the fibrillar state. Then, we demonstrated that the small molecule binds through its hydrophilic moiety to Aβ42 aggregates mainly through a hydrogen bond made by its carbonyl group. The results demonstrate that infrared nanospectroscopy enables the identification at atomic level of the interactions between a lead compound and its target, which has been to date hindered for Aβ42 aggregates due their transient and heterogeneous nature. We therefore anticipate that the use of infrared nanospectroscopy in drug discovery will open the way to systematic structure-activity relationship studies in compound optimisation campaigns.

In conclusion, this work establishes infrared nanospectroscopy as a powerful tool for characterizing interactions of drugs with protein aggregates at the single oligomer level, and identifying key interaction signatures. More broadly, the development of single molecule biophysical methodologies represents a fruitful avenue to address the challenge of unravelling the interaction of therapeutic compounds with aggregating proteins, such as small molecules, as well as antibodies. Such a molecular level understanding is key for establishing rational approaches to prevention and design pharmacological approaches to misfolding diseases.

## ONLINE METHODS

### Aβ peptides

The recombinant Aβ(M1-42) peptide (MDAEFRHDSGY EVHHQKLVFF AEDVGSNKGA IIGLMVGGVV IA), here called Aβ42, was expressed in the *E. coli* BL21 Gold (DE3) strain (Stratagene, CA, U.S.A.) and purified as described previously with slight modifications.^38^ Briefly, the purification procedure involved sonication of *E. coli* cells, dissolution of inclusion bodies in 8 M urea, and ion exchange in batch mode on diethylaminoethyl cellulose resin and lyophylization. The lyophilized fractions were further purified using Superdex 75 HR 26/60 column (GE Healthcare, Buckinghamshire, U.K.) and eluates were analyzed using SDS-PAGE for the presence of the desired protein product. The fractions containing the recombinant protein were combined, frozen using liquid nitrogen, and lyophilized again. Bexarotene was obtained from Sigma-Aldrich, while the derivative of bexarotene was obtained from ArkPharm and both were of the highest purity available.

### Sample preparation for kinetic experiments

Solutions of monomeric peptides were prepared by dissolving the lyophilized Aβ42 peptide in 6 M GuHCl. Monomeric forms were purified from potential oligomeric species and salt using a Superdex 75 10 / 300 GL column (GE Healthcare) at a flowrate of 0.5 mL/min, and were eluted in 20 mM sodium phosphate buffer, pH 8 supplemented with 200 μM EDTA and 0,02% NaN_3_. The centre of the peak was collected and the peptide concentration was determined from the absorbance of the integrated peak area using ε_280_ = 1490 l mol^-1^ cm^-1^. The obtained monomer was diluted with buffer to the desired concentration and supplemented with 20 *μ*M Thioflavin T (ThT) from a 2 mM stock. All samples were prepared in low binding eppendorf tubes on ice using careful pipetting to avoid introduction of air bubbles. Each sample was then pipetted into multiple wells of a 96-well half-area, low-binding, clear bottom and PEG coating plate (Corning 3881), 80 μL per well, in the absence and the presence of bexarotene.

### Kinetic assays

Assays were initiated by placing the 96-well plate at 37 ºC under quiescent conditions in a plate reader (Fluostar Omega, Fluostar Optima or Fluostar Galaxy, BMGLabtech, Offenburg, Germany). The ThT fluorescence was measured through the bottom of the plate with a 440 nm excitation filter and a 480 nm emission filter. The ThT fluorescence was followed for three repeats of each sample.

### Theoretical analysis

The time evolution of the total fibril mass concentration, *M(t)*, is described by the following integrated rate law:^22,42^

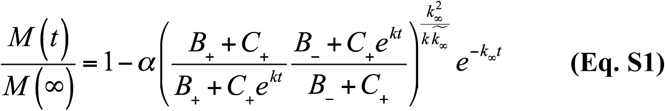

To capture the complete assembly process, only two particular combinations of the rate constants define most of the macroscopic behaviours. These are related to the rate of formation of new aggregates through primary pathways 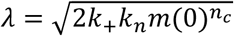 and secondary pathways 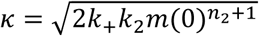, where the initial concentration of soluble monomers is denoted by *m*(0), n_c_ and n_2_ describe the dependencies of the primary and secondary pathways on the monomer concentration, and k_n_, k_+_ and k_2_ are the rate constants of the primary nucleation, elongation and secondary nucleation, respectively. Using **Eq. S1**, the experimental data of the aggregation of Aβ42 in the absence and presence of bexarotene can be described via a concomitant decrease in both *k*_*n*_ and *k*_*2*_.

### FTIR measurements

Attenuated total reflection infrared spectroscopy (ATR-FTIR) was performed using a Bruker Vertex 70 spectrometer equipped with a diamond ATR element. 2μM samples of Aβ42 aggregates in the absence and presence of 3-fold excess were centrifuged and re-suspended in buffer to a final concentration of 2 mM. Spectra were acquired with a resolution of 4 cm^-1^ and all spectra were processed using Origin Pro software. The spectra were averaged (3 spectra with 512 co-averages), smoothed applying a Savitzky-Golay filter (2^nd^ order, 9 pts).

### Atomic Force Microscopy

Atomic Force Microscopy was performed on positive functionalized mica substrates. We cleaved the mica surface and we incubated it for 1 minute with 10 µl of 0.5% (v/v) (3-Aminopropyl)triethoxysilane (APTES, from SIGMA) in Milli-Q water. Then, the substrate was rinsed three times with 1 ml of Milli-Q water and dried by gentle stream of nitrogen gas. Finally, for each sample, an aliquot of 10 µl of the solution was deposited on the functionalized surface. The droplet was incubated for 10 minutes, then rinsed by 1 ml of Milli-Q water and dried by the gentle stream of nitrogen gas. The preparation was carried out at room temperature. AFM maps were acquired by means of a NX10 (Park systems, South Korea) operating in tapping mode and equipped with a silicon tip (μmasch, 2 Nm^-1^) with a nominal radius of ∼8 nm. Image flattening and single aggregate cross-sectional dimension analysis, were performed by SPIP (Image Metrology) software.

### AFM-IR measurements, maps treatment and analysis

Analysis by nanoIR2 (Anasys Instrument, USA) was performed on atomically flat and positive gold substrates with a nominal roughness of 0.36 nm (Platypus Technologies, USA)^43^. The root mean square roughness of the AFM maps was measured by SPIP (Image metrology, Denmark). To prepare the protein samples, an aliquot of 10 µl of sample was deposited on the flat gold surface for 30 s, to reduce mass transport phenomena during drying. Successively, the droplet was rinsed by 1 ml of Milli-Q water and dried by a gentle stream of nitrogen. The morphology of the protein samples was scanned by the nanoIR microscopy system, with a rate line within 0.1-0.5 Hz and in contact mode. All AFM maps were acquired with a resolution between 2-10 nm/pixel. A silicon gold coated PR-EX-nIR2 (Anasys, USA) cantilever with a nominal radius of ∼30 nm and an elastic constant of about 0.2 N/m was used. To use gold-gold rod-like antenna effect the IR light was polarized perpendicular to the surface of deposition. The AFM images were treated and analysed using SPIP software. The height images were first and second order flattened. Spectra were collected with a laser wavelength sampling of 2 cm^-1^ with a spectral resolution of ∼1 cm^-1^ and 256 co-averages, within the range 1400-1800 cm^-1^. The spectra were acquired at a speed of 60 cm^-1^/s. All spectra and maps were acquired at the same power of background laser power, between approximately 0.35-1 mW and with a pulse width between 40-100 ns. Since the spectral background line shape slightly depends on laser power, the spectra were normalised by the QCL emission profile.

All measurements were performed at room temperature under controlled nitrogen atmosphere with residual real humidity below 5%. The spectra were acquired by using measuring off-resonance, on the left of the IR amplitude maximum resonant frequency.^35,44,45^

### Spectra analysis

Spectra were analysed using the microscope’s built-in Analysis Studio (Anasys) and OriginPRO. The spectra where despiked, an adjacent averaging filter (5pts) and a Savitzky-Golay filter (15 pts, 2^nd^ order) were applied in series and then the spectra were normalised to one respect to the maximum of intensity. The structural contributions were investigated by second derivative analysis to deconvolution of the amide band I.^24,26,46^ The second derivatives were smoothed by a Savitzky-Golay filter (2^nd^ order, 7 pts).

For the PCA, the spectra were smoothed and normalised spectra, as described above, and further baselined to zero in the range 1400-1800 cm^-1^. PCA was performed by means of OriginPRO by considering a mean-centered covariance matrix. In all cases the first three PCs accounted for more than 85% of spectral variance.

## ACKNOWLEDGEMENTS

### FUNDING

We thank Darwin College and Swiss National Fondation for Science (SNF) for the financial support (grant number P2ELP2_162116 and P300P2_171219). The research leading to these results has received funding from the Wellcome Trust and the European Research Council under the European Union’s Seventh Framework Programme (FP7/2007-2013) through the ERC grant PhysProt (agreement n° 337969).

### AUTHORS CONTRIBUTIONS

F.S.R., J.H. conceived the project. F.S.R., J. H. and S.C. performed the experiments. F.S.R. analysed the data. F.S.R., M.V. and T.P.J.K. wrote and commented the article.

### COMPETING INTERESTS

The authors declare no competing interests.

### DATA

All data needed to evaluate the conclusions in the paper are present in the paper and/or the Supplementary Materials. Additional data are available from authors upon request.

### CORRESPONDING AUTHOR

Correspondence should be addressed to: F.S. Ruggeri, Michele Vendruscolo and Tuomas P. J. Knowles.

## Supplementary Materials

### SUPPLEMENTARY FIGURES

**Figure S1.**
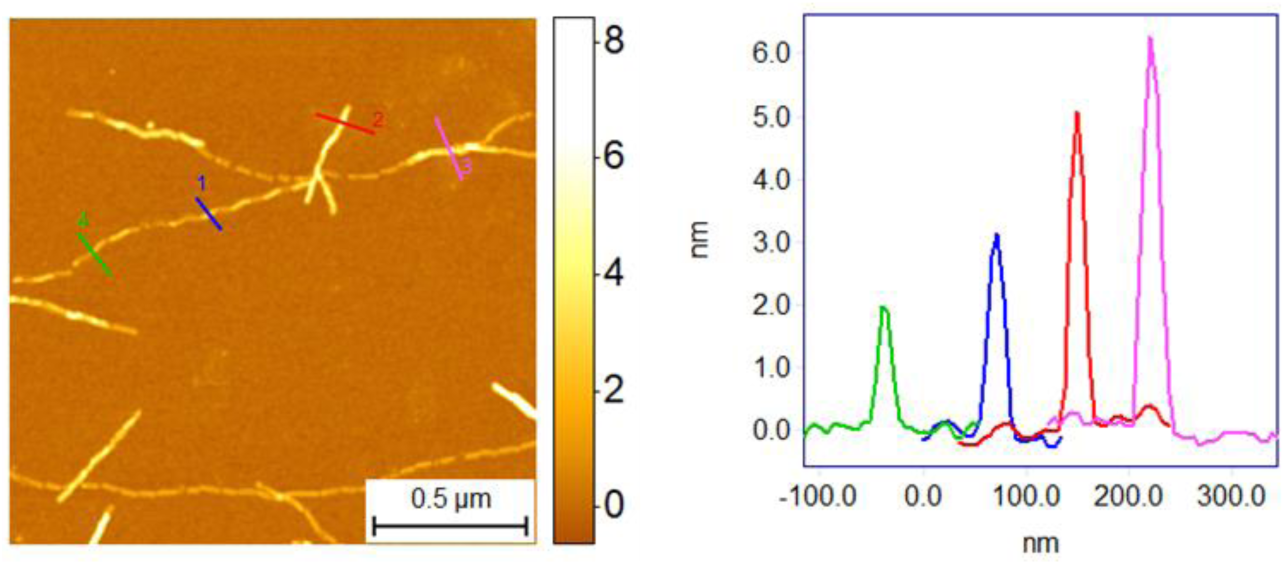
Cross-sectional dimensions of Aβ42 fibrils on mica substrate. a) High resolution 3-D morphology map. b) Cross-sectional height if the aggregates.

**Figure S2.**
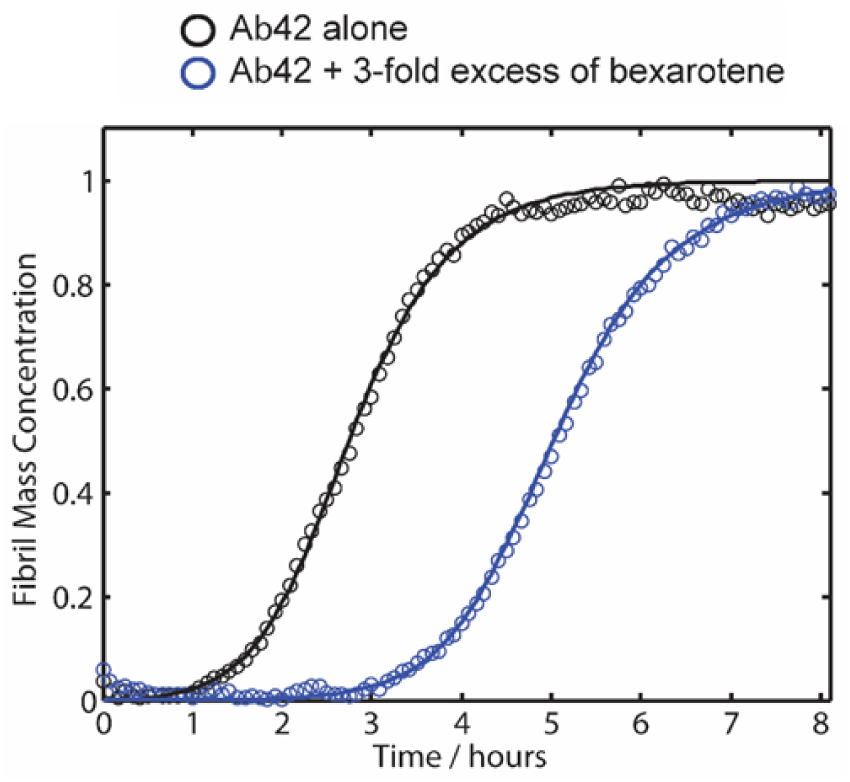
Kinetics of aggregation of Aβ42 in presence of bexarotene. Kinetic profiles of the aggregation reaction of 2 μM Aβ42 in the absence or in the presence of a 3:1 concentration ratio of bexarotene to Aβ42. The solid lines show the best fit of the experimental data (open circles) when primary and secondary pathways are both inhibited by bexarotene.

**Figure S3.**
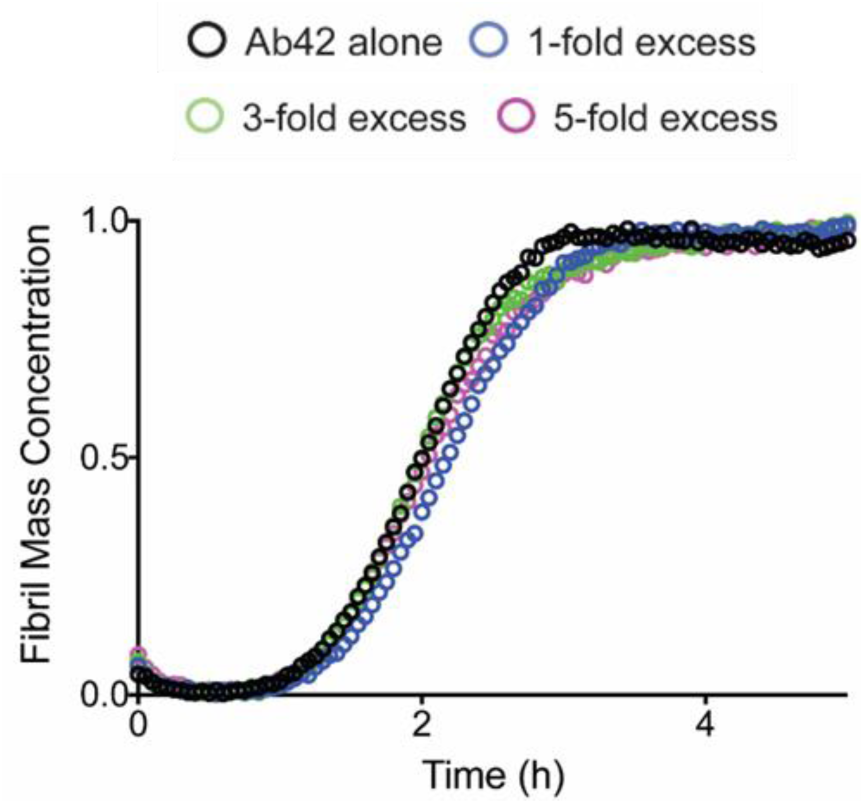
Kinetics of aggregation of Aβ42 in presence of the derivative of bexarotene. Kinetic profiles of the aggregation reaction of 2 μM Aβ42 in the absence or in the presence of 1-fold to 5-fold excess of a derivative of bexarotene. Note that no significant delay in the aggregation is observed in the presence of this compound.

**Figure S4.**
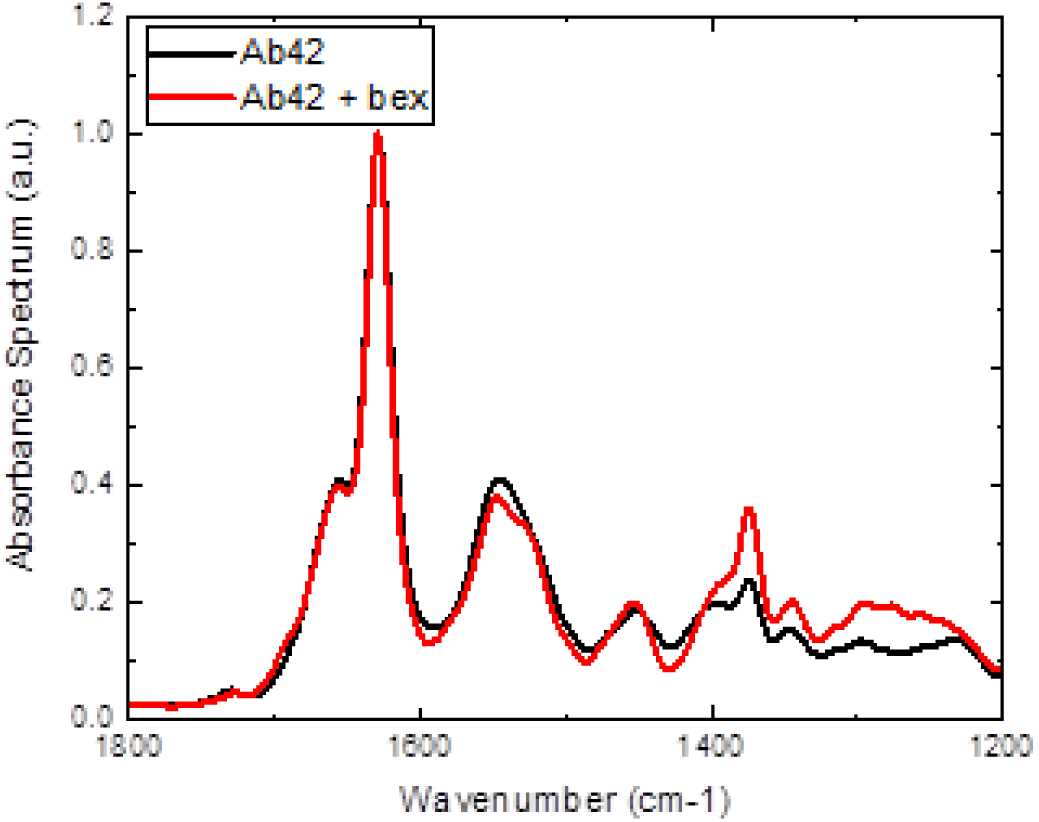
ATR spectroscopy of Aβ42 fibrillar aggregates in absence and presence of bexarotene. The sample is prepared at a concentration of Aβ42 of 2 mM, which is far beyond the capabilities of conventional bulk FTIR. Thus, we acquired the spectra in air conditions, which did not allow to observe any difference between the aggregates with and without the small molecule.

**Figure S5.**
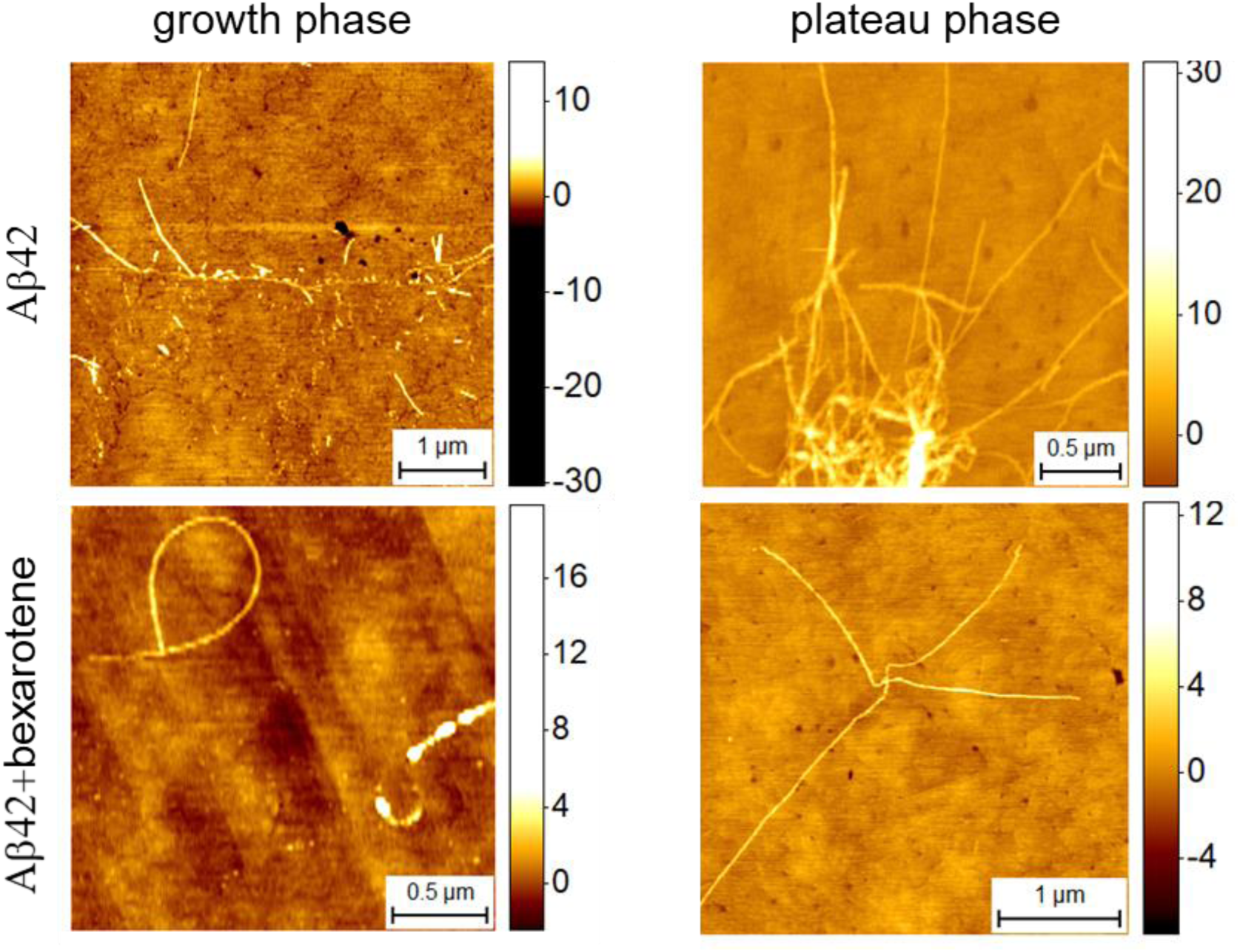
Morphology of Aβ42 aggregates in the growth phase and plateau phase of aggregation on gold substrates.

**Figure S6.**
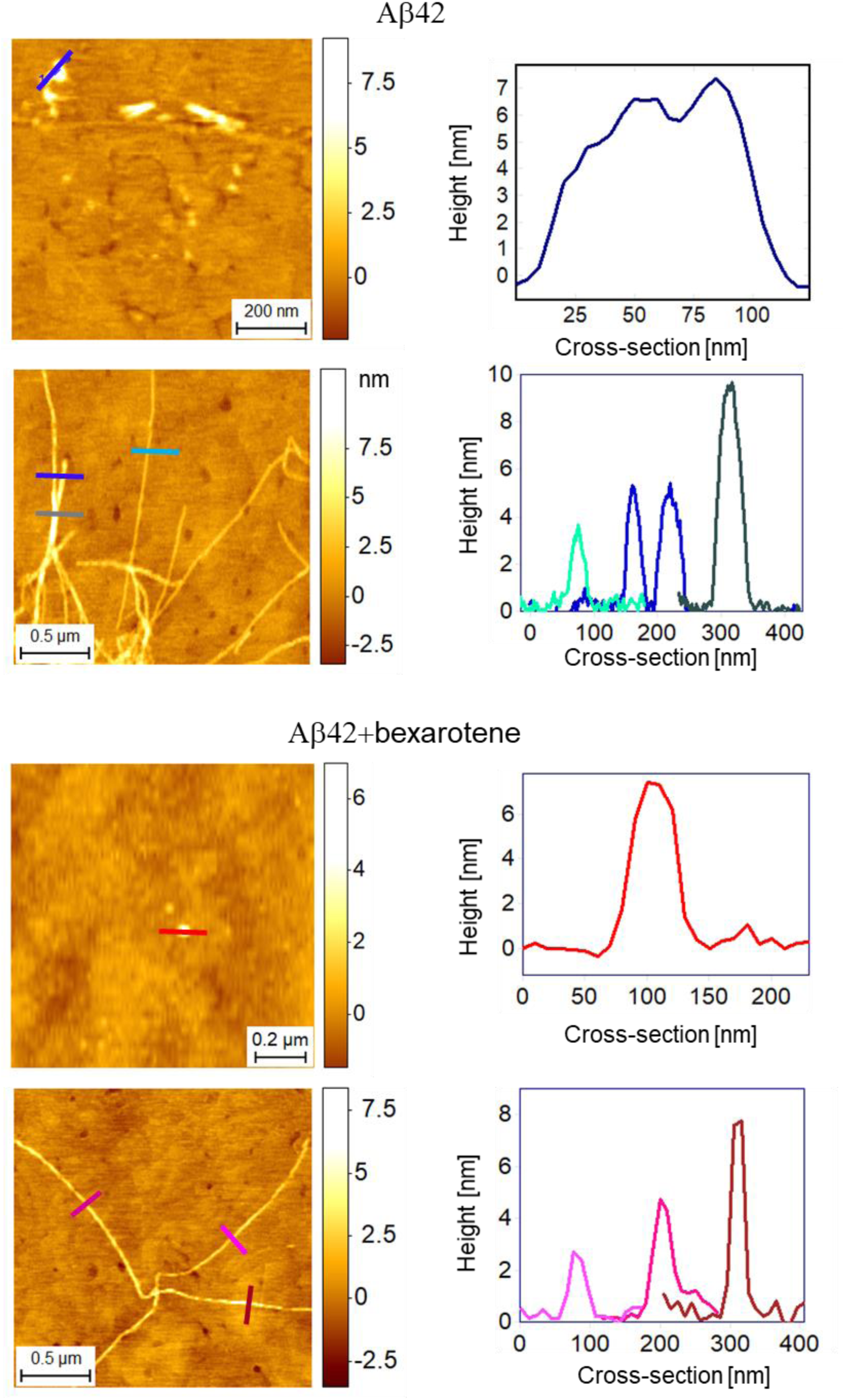
Cross-sectional dimensions of Aβ42 aggregates incubated in presence and absence of bexarotene on gold substrates and where the AFM-IR spectra were acquired.

**Figure S7.**
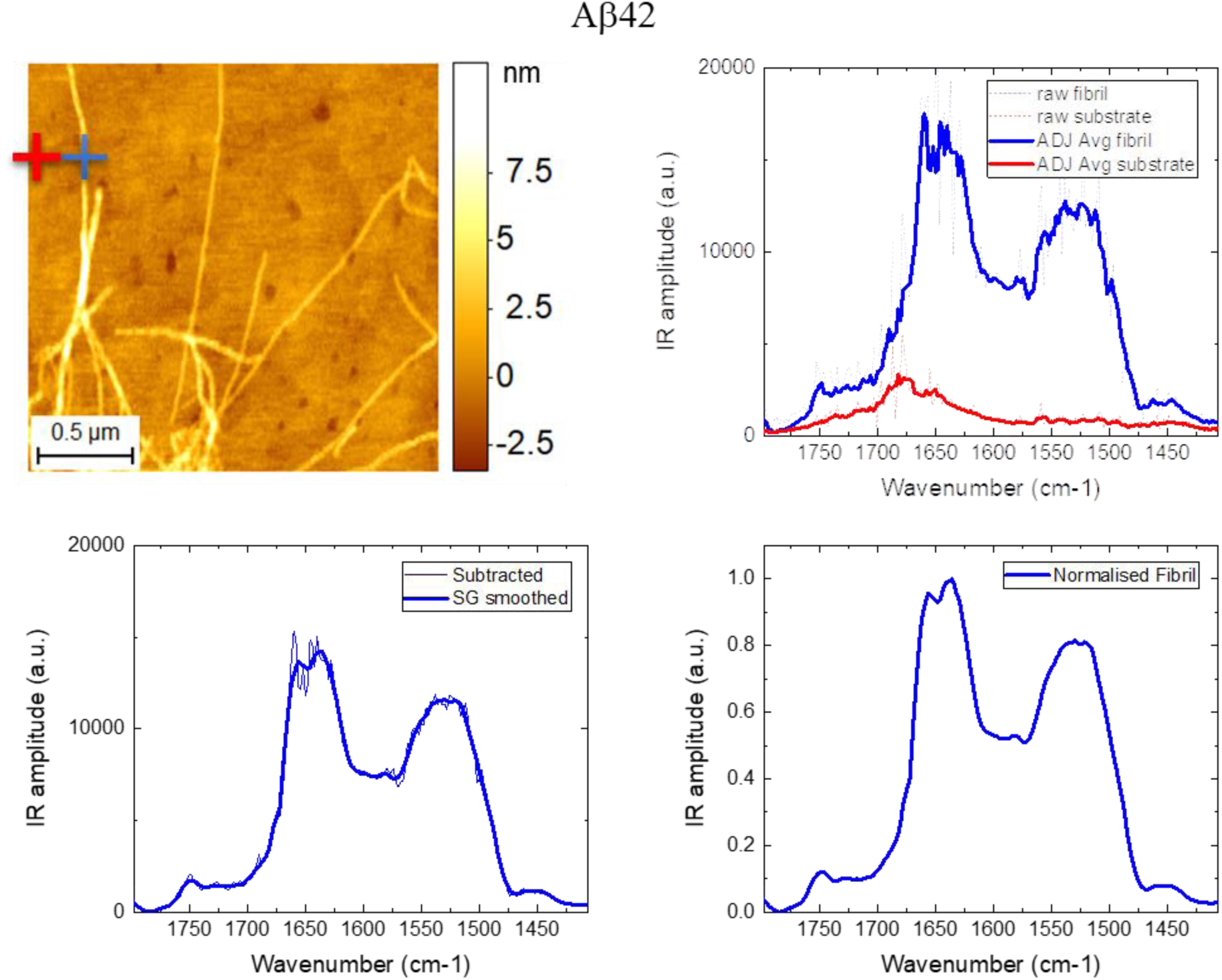
Acquisition and post-processing of AFM-IR spectra. a-b) Nanoscale IR spectra are acquire on the top of the aggregate of interest (blue cross) and on the surrounding substrate (red cross). c) The two spectra are subtracted to remove tip contamination and residual substrate absorbance from the spectrum of the aggregate, which is then smoothed by a Savitzky-Golay (15 pts, 2^nd^ order) filter and then d) normalised to the maximum of absorption.

**Figure S8.**
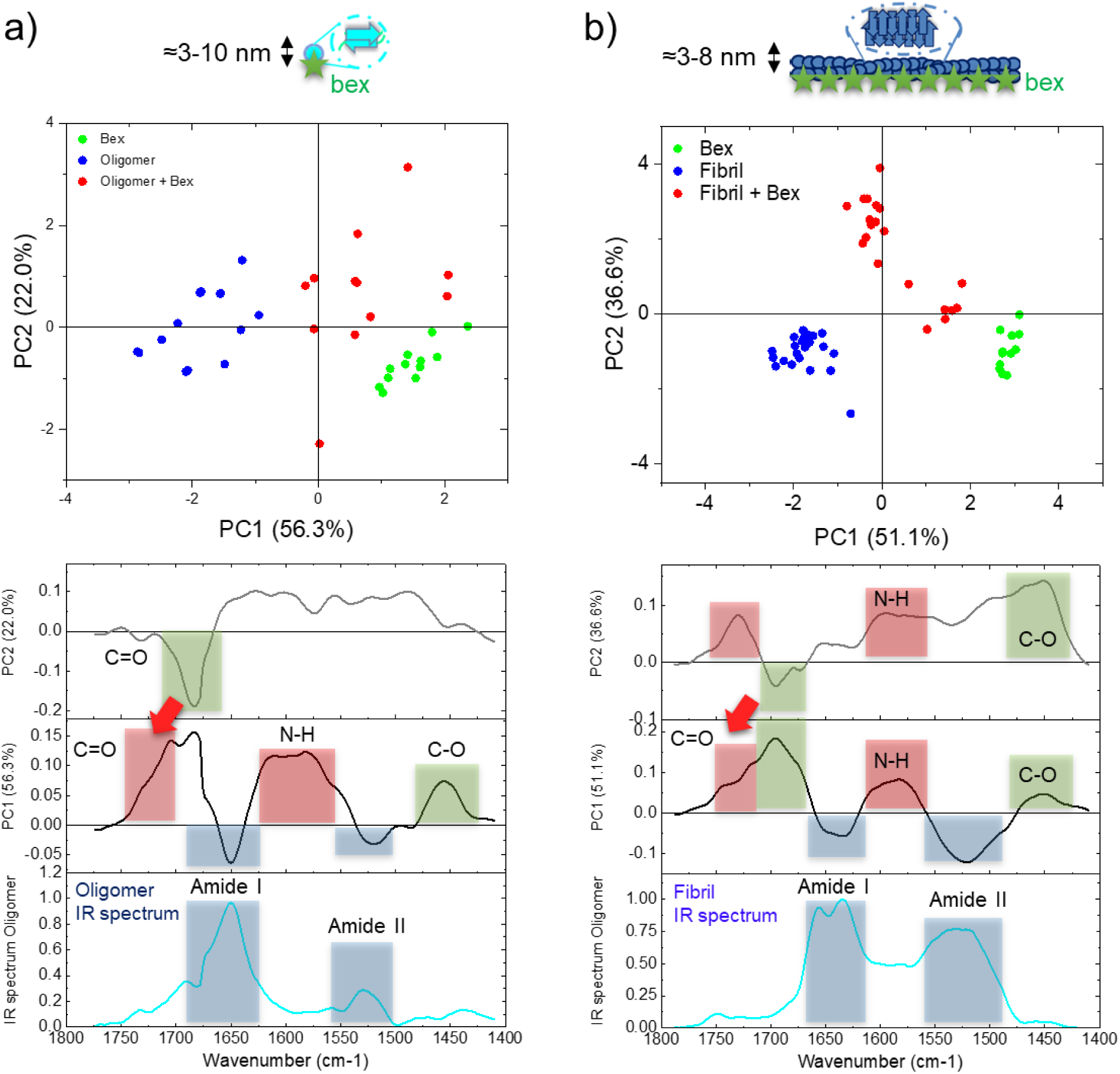
PCA analysis on oligomeric and fibrillar aggregates. Score plot and relative loading plots of the analysis on a) oligomeric species and bexarotene, b) fibrillar species and bexarotene.

**Figure S9.**
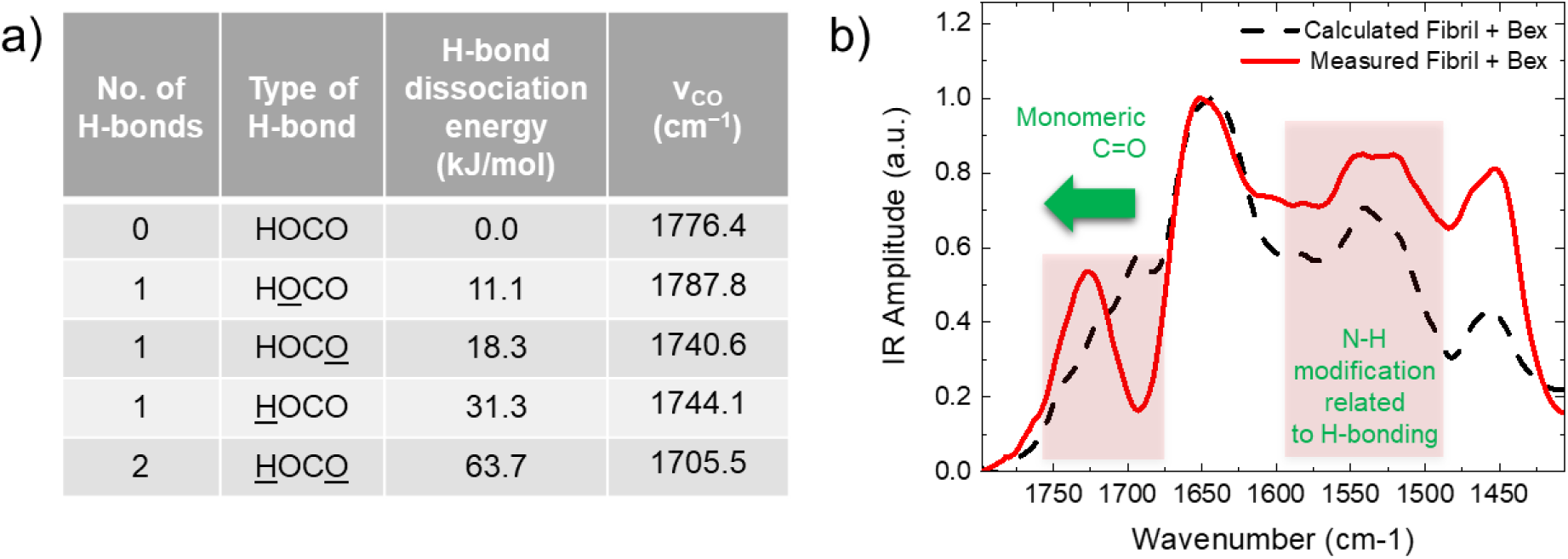
Bexarotene interacts with Aβ42 aggregates through hydrogen bonding. (a) Energies and wavenumbers characteristic of different coordination states for the carboxylic group, adapted from ^41^. If we consider the bidentate and the single acylic coordination’s (rows in yellow), we see that the IR band shifts from 1705.5 cm^−1^ to 1740.6 cm^−1^, passing from the former to the latter. (b) The same behaviour is observed in our experimental results where it is evident the shift of the CO stretching from ∼1700 cm^−1^ to ∼1730 cm^−1^ from the spectrum of bexarotene alone to the one of the aggregates incubated with the molecule. b) To further stress this result, we have calculated the sum of the measured spectra of bexarotene and of the amyloid aggregates alone. This calculated spectrum differs significantly from the spectrum measured on the aggregates incubated with the small molecule in the region of the C=O and N-H stretching. The comparison of the experimental measured spectrum and of the calculated spectrum summing the experimental IR signatures of fibrils and bexarotene alone in its dimeric form confirms that the only differences in the spectra are related to a monomeric hydrogen-bonding.

## REFERENCES

1 Alzheimer’s, A. 2012 Alzheimer’s disease facts and figures. Alzheimers Dement 8, 131–168, (2012).

2 Knowles, T. P. J., Vendruscolo, M. & Dobson, C. M. The amyloid state and its association with protein misfolding diseases. Nat Rev Mol Cell Biol 15, 384–396, (2014).

3 Selkoe, D. J. Alzheimer’s Disease: Genes, Proteins, and Therapy. Physiological Reviews 81, 741–766, (2001).

4 Haass, C., x & Selkoe, D. J. Soluble protein oligomers in neurodegeneration: lessons from the Alzheimer’s amyloid β-peptide. Nature Reviews Molecular Cell Biology 8, 101–112, (2007).

5 Dobson, C. M. Protein folding and misfolding. Nature 426, 884–890, (2003).

6 Bieschke, J. Natural compounds may open new routes to treatment of amyloid diseases. Neurotherapeutics 10, 429–439, (2013).

7 Chen, J. X. & Yan, S. S. Role of mitochondrial amyloid-beta in Alzheimer’s disease. J Alzheimers Dis 20 Suppl 2, S569–578, (2010).

8 Kroth, H. et al. Discovery and Structure Activity Relationship of Small Molecule Inhibitors of Toxic beta-Amyloid-42 Fibril Formation. Journal of Biological Chemistry 287, 34786–34800, (2012).

9 Lansbury, P. T. & Lashuel, H. A. A century-old debate on protein aggregation and neurodegeneration enters the clinic. Nature 443, 774–779, (2006).

10 Nie, Q., Du, X. G. & Geng, M. Y. Small molecule inhibitors of amyloid beta peptide aggregation as a potential therapeutic strategy for Alzheimer’s disease. Acta Pharmacologica Sinica 32, 545–551, (2011).

11 Porat, Y., Abramowitz, A. & Gazit, E. Inhibition of amyloid fibril formation by polyphenols: Structural similarity and aromatic interactions as a common inhibition mechanism. Chemical Biology & Drug Design 67, 27–37, (2006).

12 Sinha, S. et al. Lysine-Specific Molecular Tweezers Are Broad-Spectrum Inhibitors of Assembly and Toxicity of Amyloid Proteins. Journal of the American Chemical Society 133, 16958–16969, (2011).

13 Cummings, J. & Zhong, K. Biomarker-Driven Therapeutic Management of Alzheimer’s Disease: Establishing the Foundations. Clinical Pharmacology & Therapeutics 95, 67–77, (2014).

14 Selkoe, D. J. Resolving controversies on the path to Alzheimer’s therapeutics. Nature Medicine 17, 1060–1065, (2011).

15 Arosio, P., Vendruscolo, M., Dobson, C. M. & Knowles, T. P. J. Chemical kinetics for drug discovery to combat protein aggregation diseases. Trends in Pharmacological Sciences 35, 127–135, (2014).

16 Butterfield, S. & Lashuel, H. Amyloidogenic Protein-Membrane Interactions: Mechanistic Insight from Model Systems. Angewandte Chemie International Edition 49, 5628–5654, (2010).

17 Campioni, S. et al. A causative link between the structure of aberrant protein oligomers and their toxicity. Nature Chemical Biology 6, 140–147, (2010).

18 Mannini, B. et al. Toxicity of Protein Oligomers Is Rationalized by a Function Combining Size and Surface Hydrophobicity. ACS Chemical Biology 9, 2309–2317, (2014).

19 Shankar, G. M. et al. Amyloid-beta protein dimers isolated directly from Alzheimer’s brains impair synaptic plasticity and memory. Nature Medicine 14, 837–842, (2008).

20 Walsh, P., Neudecker, P. & Sharpe, S. Structural properties and dynamic behavior of nonfibrillar oligomers formed by PrP(106-126). Journal of the American Chemical Society 132, 7684–7695, (2010).

21 Habchi, J. et al. An anticancer drug suppresses the primary nucleation reaction that initiates the production of the toxic Aβ42 aggregates linked with Alzheimer’s disease. Science Advances 2, (2016).

22 Cohen, S. I. a. et al. Proliferation of amyloid-β42 aggregates occurs through a secondary nucleation mechanism. Proceedings of the National Academy of Sciences of the United States of America 110, 9758–9763, (2013).

23 Scheidt, T. et al. Secondary nucleation and elongation occur at different sites on Alzheimer’s amyloid-β aggregates. Science Advances 5, eaau3112, (2019).

24 Ruggeri, F. S. et al. Infrared Nanospectroscopy Characterization of Oligomeric and Fibrillar Aggregates during Amyloid Formation. Nat Commun 6, 7831 (2015). <https://www.ncbi.nlm.nih.gov/pubmed/26215704>.

25 Muller, T. et al. Nanoscale spatially resolved infrared spectra from single microdroplets. Lab Chip 14, 1315–1319, (2014).

26 Qamar, S. et al. FUS Phase Separation Is Modulated by a Molecular Chaperone and Methylation of Arginine Cation-π Interactions. Cell 173, 720–734, (2018).

27 Ruggeri, F. S. et al. Identification of Oxidative Stress in Red Blood Cells with Nanoscale Chemical Resolution by Infrared Nanospectroscopy. Int J Mol Sci 19, 2582, (2018).

28 Volpatti, L. R. et al. Micro- and Nanoscale Hierarchical Structure of Core-Shell Protein Microgels. Journal of Materials Chemistry B 4, 7989–7999, (2016).

29 Ruggeri, F. S. et al. Nanoscale Studies Link Amyloid Maturity with Polyglutamine Diseases Onset. Sci Rep 6, 31155, (2016).

30 Ruggeri, F. S. et al. Concentration-dependent and surface-assisted self-assembly properties of a bioactive estrogen receptor alpha-derived peptide. Journal of peptide science : an official publication of the European Peptide Society 21, 95–104, (2015).

31 Dazzi, A. & Prater, C. B. AFM-IR: Technology and Applications in Nanoscale Infrared Spectroscopy and Chemical Imaging. Chem Rev 117, 5146–5173, (2017).

32 Centrone, A. in Annual Review of Analytical Chemistry, Vol 8 Vol. 8 Annual Review of Analytical Chemistry (eds R. G. Cooks & J. E. Pemberton) 101-126 (Annual Reviews, 2015).

33 Galante, D. et al. A critical concentration of N-terminal pyroglutamylated amyloid beta drives the misfolding of Ab1-42 into more toxic aggregates. Int J Biochem Cell Biol 79, 261–270, (2016).

34 Ruggeri, F. S., Habchi, J., Cerreta, A. & Dietler, G. AFM-Based Single Molecule Techniques: Unraveling the Amyloid Pathogenic Species. Current pharmaceutical design 22, 3950–3970, (2016).

35 Ramer, G., Ruggeri, F. S., Levin, A., Knowles, T. P. J. & Centrone, A. Determination of Polypeptide Conformation with Nanoscale Resolution in Water. ACS Nano 12, 6612–6619, (2018).

36 Lu, F., Jin, M. Z. & Belkin, M. A. Tip-enhanced infrared nanospectroscopy via molecular expansion force detection. Nature Photonics 8, 307–312, (2014).

37 Ruggeri, F. S., Mannini, B., Schmid, R., Vendruscolo, M. & Knowles, T. P. J. Single molecule secondary structure determination of proteins through infrared absorption nanospectroscopy. Nature Communications 11, 2945, (2020).

38 Hellstrand, E., Boland, B., Walsh, D. M. & Linse, S. Amyloid beta-Protein Aggregation Produces Highly Reproducible Kinetic Data and Occurs by a Two-Phase Process. Acs Chemical Neuroscience 1, 13–18, (2010).

39 Michaels, T. C. T. et al. Dynamics of oligomer populations formed during the aggregation of Alzheimer’s Aβ42 peptide. Nature Chemistry 12, 445–451, (2020).

40 Abrosimova, K. V., Shulenina, O. V. & Paston, S. V. FTIR study of secondary structure of bovine serum albumin and ovalbumin. Journal of Physics: Conference Series 769, 12016, (2016).

41 Nie, B., Stutzman, J. & Xie, A. A Vibrational Spectral Maker for Probing the Hydrogen-Bonding Status of Protonated Asp and Glu Residues. Biophys J 88, 2833–2847, (2005).

42 Cohen, S. I. A., Vendruscolo, M., Dobson, C. M. & Knowles, T. P. J. From macroscopic measurements to microscopic mechanisms of protein aggregation. Journal of molecular biology 421, 160–171, (2012).

43 Miller, M. S., Ferrato, M.-A., Niec, A., Biesinger, M. C. & Carmichael, T. B. Ultrasmooth Gold Surfaces Prepared by Chemical Mechanical Polishing for Applications in Nanoscience. Langmuir 30, 14171–14178, (2014).

44 Ruggeri, F. S., Sneideris, T., Chia, S., Vendruscolo, M. & Knowles, T. P. J. Characterizing Individual Protein Aggregates by Infrared Nanospectroscopy and Atomic Force Microscopy. J Vis Exp, (2019).

45 Ramer, G., Reisenbauer, F., Steindl, B., Tomischko, W. & Lendl, B. Implementation of Resonance Tracking for Assuring Reliability in Resonance Enhanced Photothermal Infrared Spectroscopy and Imaging. Applied Spectroscopy 71, 2013–2020, (2017).

46 Shimanovich, U. et al. Silk Micrococoons for Protein Stabilisation and Molecular Encapsulation. Nat Commun 8, 15902 (2017). <https://www.ncbi.nlm.nih.gov/pubmed/28722016>.

